# Pupil dilation is sensitive to semantic ambiguity and acoustic degradation

**DOI:** 10.1101/2020.02.19.955609

**Authors:** Mason Kadem, Björn Herrmann, Jennifer M. Rodd, Ingrid S. Johnsrude

## Abstract

Speech comprehension is often challenged by background noise or other acoustic interference. It can also be challenged by linguistic factors, such as complex syntax, or the presence of words with more than one meaning. Pupillometry is increasingly recognized as a technique that provides a window onto acoustic challenges, but this work has not been well integrated with an older literature linking pupil dilation to “mental effort”, which would include linguistic challenges. Here, we measured pupil dilation while listeners heard spoken sentences with clear sentence-level meaning that contained words with more than one meaning (“The shell was fired towards the tank”) or matched sentences without ambiguous words (“Her secrets were written in her diary”). This semantic-ambiguity manipulation was crossed with an acoustic manipulation: two levels of a 30-talker babble masker in Experiment 1; and presence or absence of a pink noise masker in Experiment 2. Speech comprehension, indexed by a semantic relatedness task, was high (above 82% correct) in all conditions. Pupils dilated when sentences included semantically ambiguous words compared to matched sentences and when maskers were present compared to absent (Experiment 2) or were more compared to less intense (Experiment 1). The current results reinforce the idea that many different challenges to speech comprehension, that afford different cognitive processes and are met by the brain in different ways, manifest as an increase in pupil dilation.

## Introduction

Following and understanding one particular conversational partner, despite interference from other sources, is a feat most of us accomplish effortlessly every day. However, many processes are required to analyze a complex auditory signal, consisting of many different sound sources, so that one source (i.e., a voice) can be identified, tracked, and understood. The process is complicated by the enormous variability of speech – speech is often in an unfamiliar accent or voice, distorted or degraded, or masked by other sounds. Different acoustic challenges may depend on different cognitive resources for speech comprehension to be successful. For example, competing voices may be acoustically different enough from the target speech signal that the target can be isolated and tracked: this requires cognitive control and distracter suppression (Johnsrude and Rodd, 2016), for perceptually similar (speech-like) maskers to not be mistaken for the target. In contrast, when speech is masked energetically (i.e., by a noise, with an overlapping frequency spectrum) some of the speech signal is obliterated and missing information must be inferred from the bits of speech that are perceived. This probably requires effective working memory and access to semantic knowledge (Johnsrude and Rodd, 2016).

Linguistic factors also challenge speech comprehension (Gibson, 1998; Gibson and Pearlmutter, 1998). Sometimes utterances are simple and straightforward, such as the statement “The dog barked at the squirrel”, but other times, the linguistic structure is more complex (“It was the squirrel at which the dog barked”), or the utterance lacks clear (to the listener) meaningfulness at the word and/or sentence level that would aid comprehension, because words have multiple meanings, or are uncommon (“The bark ruffled the sciurid”). Again, the cognitive resources recruited to compensate for such linguistic demands probably differ, depending on the demand (Gibson, 1998; Johnsrude and Rodd, 2016; Van Hedger and Johnsrude, in press).

Speech understanding can be particularly challenging for those with hearing loss. Substantially greater demands must be placed on knowledge-based, compensatory mechanisms in hearing-impaired individuals, who report listening in such situations to be effortful (Nachtegaal et al., 2009; Hornsby, 2013). This listening effort is a serious obstacle to communication, affecting all aspects of a person’s life (Banh et al., 2012; Pichora-Fuller et al., 2016). Listening effort is therefore increasingly recognized as a useful concept to understand the hearing problems many normally aging adults experience in their everyday lives (Eckert et al., 2016; Johnsrude and Rodd, 2016; Lemke and Besser, 2016; Pichora-Fuller et al., 2016; Strauss and Francis, 2017; Peelle, 2018; Winn et al., 2018).

Listening effort may explain variance in behavior that is not captured by standard hearing assessment (e.g., audiometry). Measuring listening effort effectively has thus become a major endeavor in the hearing science and audiology communities.

Subjective ratings are a common way to assess listening effort (Gatehouse and Noble, 2004; Larsby et al., 2005; Wendt et al., 2016; Alhanbali et al., 2017; Krueger et al., 2017). However, subjective measures have a host of limitations such as context effects (participants may rate their experienced effort relative to different conditions within an experiment rather than in absolute terms of their experience) and inter-subject differences in scale use. Moreover, established scales are only appropriate for use with older children and adults; animals and babies cannot do the task, and ‘effort’ may be conceptualized differently in different cultures, limiting comparative research. Objective, physiological measures can also provide a window onto listening effort. Pupillometry – the measurement of the dilation of an individual’s pupil, has long been used to study “mental effort” (Kahneman and Beatty, 1966; Beatty, 1982; Sirois and Brisson, 2014). This approach has, more recently, sparked great interest among hearing scientists and audiologists because of its potential applicability in the clinic (Winn et al., 2018; Zekveld et al., 2018).

Pupillometry studies focusing on acoustic challenges during listening demonstrate that the pupil is typically larger when individuals listen to acoustically degraded speech compared to acoustically less degraded speech (Zekveld et al., 2010; Zekveld et al., 2014; Winn et al., 2015; Wendt et al., 2016; Miles et al., 2017; Borghini and Hazan, 2018), although pupil dilation may saturate for highly degraded, but intelligible speech signals (Zekveld and Kramer, 2014; Ohlenforst et al., 2017).

Pupillometric measurements of cognition have a long history and we have long known that any challenge that increases the brain’s processing load, will dilate the pupil (Kahneman and Beatty, 1966; Kahneman, 1973), but pupillometry has not been used very often to study the effects of linguistic challenges on speech comprehension. Two studies have shown that pupil dilation is enhanced for syntactically complex, object-first sentences compared to less complex, subject-first sentences (Wendt et al., 2016; Ayasse and Wingfield, 2018), indicating that pupillometry can provide a window onto linguistic challenges during speech comprehension.

The effect of semantic ambiguity on pupil dilation during sentence comprehension is less clear, although other work suggests that the presence of semantically ambiguous words is cognitively demanding (Rodd et al., 2005; Rodd et al., 2010a; Johnsrude and Rodd, 2016; Rodd, in press). Indeed, isolated words that are semantically difficult to process (based on word frequency, familiarity, and others) (Chapman and Hallowell, 2015) or words presented under lexical competition (Kuchinsky et al., 2013) lead to larger pupil dilation compared to words that are semantically easier to process. Moreover, sentences with weak semantic constraints have been shown to lead to larger pupil dilation compared to sentences with strong semantic constraints (Winn, 2016). However, sentences whose meaning is unambiguous, but which contain multiple ambiguous words (e.g., The **shell** was **fired** towards the **tank**) are common in real life, but it is unknown whether pupillometry is sensitive to the demands imposed by such sentences.

Acoustic and linguistic challenges may interact in their effect on pupil dilation: The effect of linguistic challenges may be particularly prominent under high compared to low acoustic challenges (Kuchinsky et al., 2013; Wendt et al., 2016). In contrast, high cognitive load my saturate pupil dilation (Zekveld and Kramer, 2014; Ohlenforst et al., 2017), such that acoustic and linguistic challenges may be sub-additive in their effects on pupil dilation.

In the current study, we conducted two experiments to investigate whether semantic ambiguity and speech clarity affect sentence comprehension and pupil dilation. In both experiments, we presented semantically ambiguous sentences containing words with more than one meaning such as “the **shell** was **fired** towards the **tank**” and semantically less ambiguous control sentences (Rodd et al., 2010a). In Experiment 1, sentences were presented in an ongoing multi-talker background noise either under a high speech-to-noise ratio (low demands) or a low speech-to-noise ratio (high demands). In Experiment 2, speech clarity was manipulated by adding a meaningless pink noise whose energy was perfectly correlated with a sentence’s amplitude envelope and which effectively maintained a similar level of acoustic degradation throughout a sentence. We expected that pupil dilation would be increased for acoustically and semantically challenging sentences compared to less challenging ones. We expected that acoustic and linguistic challenges may interact in their effect on pupil dilation.

## Methods and Materials

Data are publicly available at https://osf.io/9kfn4/

### Participants

Seventy-three graduate and undergraduate students from The University of Western Ontario (Canada) were recruited in two experiments (Experiment 1: N=38, mean age: 20.4 years, range: 18-33 years, 26 females; Experiment 2: N=35, mean age: 19 years, range: 17-21 years, 15 females). One person who participated in Experiment 1 did not provide information regarding age and sex, but was recruited from the same student population. Data from one additional participant recorded for Experiment 2 were excluded due to failure in data storage. Participants self-reported having normal hearing, normal or corrected-to-normal vision, and no neurological disorders in their history. Participants gave written informed consent and received course credits or were paid $10 per hour for their participation. The experimental protocols were approved by the Research Ethics Board of the University of Western Ontario (protocol ID: HSREB 106570) and are in line with the declaration of Helsinki.

### Auditory stimuli and task

We utilized sentence materials from previous studies, in which the effect of sentence ambiguity on behavior and on brain activity were investigated (Rodd et al., 2005; Rodd et al., 2010a). Two conditions were utilized. In the high-ambiguity (HA) condition, sentences contained two or more ambiguous words (e.g., The **shell** was **fired** towards the **tank**) but the sentence meaning was not ambiguous. Sentences in the low-ambiguity (LA) condition contained no highly ambiguous words (e.g., Her secrets were written in her diary). The 118 (59 HA and 59 LA) original sentences were in British English and they were re-recorded by a female English speaker native to southern Ontario Canada. The duration of sentences ranged from 1.4 s to 4.8 s. The HA and LA sentences were matched on duration and psycholinguistic parameters (words, imageability, concreteness, and word frequency; Rodd et al., 2005).

In Experiment 1 (Figure 1A), background noise was added to sentences either at a low or at a high speech-to-noise (SNR) ratio. Background noise in Experiment 1 was a 30-talker babble with a longterm frequency spectrum of the current sentence materials and a flat amplitude envelope. That is, the current set of sentences was concatenated 30 times in random order, and then averaged across the 30 streams (Wagner et al., 2003). The 30-talker babble was cut and added to target sentences such that the babble noise started three seconds before sentence onset and ended 1.2 s after sentence offset. Starting the noise prior to sentence onset effectively provided acoustic cues to help segregation of speech and noise. Since the envelope of the 30-talker babble was flat, whereas the amplitude envelope of speech fluctuated naturally, masking was not constant throughout a sentence, but varied with the energy in the speech signal. The noise level was the same for all conditions (which avoided providing cues at the beginning of a trial as to which condition is presented), whereas the level of the sentence was adjusted to a signal-to-noise ratio of +6 dB (high SNR) or to a signal-to-noise-ratio of 0 dB (low SNR).

**Figure 1.**
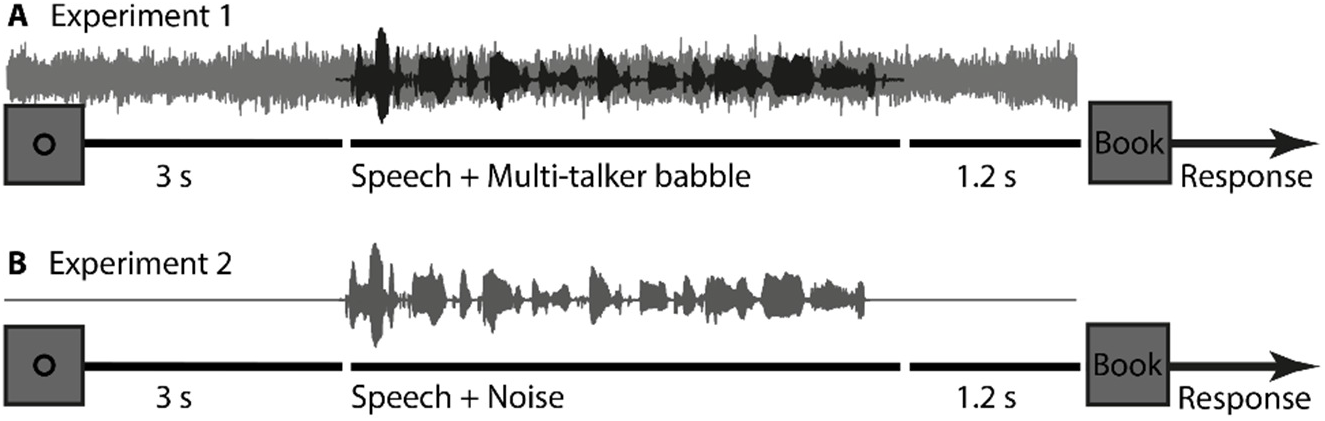
Experimental designs for Experiments 1 and 2. Schematic timeline of a trial in Experiment 1 (A) and Experiment 2 (B). A trial started three seconds prior to sentence onset with a visual fixation ring (and in Experiment 1 with the onset of the background babble noise). 1.2 s after sentence offset a probe word was presented visually. Participants were asked to indicate whether the probe word was semantically related or unrelated to the sentence.

In Experiment 2 (Figure 1B), sentences were either presented under clear conditions or with added background noise. The background noise was created uniquely for each sentence by applying the amplitude envelope of the target sentence on that trial to pink noise (1/f noise) using the Hilbert transform (30-Hz low-pass filtered; Butterworth). The original sentence and the sentence-specific modulated pink noise were added at a signal-to-noise ratio of −2 dB SNR. Since the signal and masker had the same envelope, the masking level, and thus acoustic degradation, was constant over the period of the sentence. All stimuli (including clear and those with noise added) were matched in their root-mean-square intensity level.

Both experiments were 2 × 2 factorial within-subject designs (Clarity × Ambiguity [LA, HA]). For each participant, 56 LA and 56 HA sentences were randomly selected from the 59 that were available. Half of the LA (N=28) and HA (N=28) sentences were randomly assigned to the low SNR condition (Experiment 1: 0 dB SNR babble; Experiment 2: −2 dB SNR pink noise), whereas the other half of the LA and HA sentences was assigned to the high SNR condition (Experiment 1: +6 dB SNR babble; Experiment 2: clear). Randomization was unique for each participant. In each experiment, sentences in the four conditions were presented in four blocks. Seven sentences per condition were presented within each block (N=28 trials per block), for a total of 112 (56 HA and 56 LA) sentences per person. Sentences were presented pseudo-randomly such that no more than three sentences of the same ambiguity level and two sentences of the same clarity level could occur in a row. Each participant heard each sentence only once.

### Procedure and data recording

Participants were tested in a dim, quiet room while wearing headphones (Sennheiser HD 25-SP II). Sentences were presented via a Steinberg UR22 (Steinberg Media Technologies) external sound card. Experimental procedures were controlled using Psychtoolbox in MATLAB (v2015b, Mathworks Inc.). Prior to the main experimental procedures, the hearing threshold was determined for each participant using a method-of-limits procedure described in detail in our previous work (Herrmann and Johnsrude, 2018). This procedure entailed alternating trials of progressively increasing or decreasing 12-second long pink noise over time by 5.4 dB/s. Participants indicated when they could no longer hear the noise (progressively decreasing intensity trial) or when they started to hear the noise (progressively increasing intensity trial). Each of the progressively increasing and decreasing intensity trials were presented six times, and at the time of the button press, the corresponding sound intensity during a trial was collected. Finally, the intensities from the twelve trials were averaged to determine the individual 50% hearing threshold. In both experiments, sounds were presented at 45 dB above the individual’s threshold (sensation level).

During the experiments, participants rested their head on a chin and forehead rest (EyeLink 1000 Tower mount) facing a screen at a distance of 670 mm. Pupil area and eye movements were recorded continuously from the left eye using an integrated infrared camera (eye tracker 1000; SMI, Needham, MA) at a sampling rate of 500 Hz. Nine-point fixation was used for eye-tracker calibration (McIntire et al., 2014).

During the experiments, each trial was structured as follows. Presentation of a fixation ring (black on grey [100 100 100] RGB background) started three seconds before sentence onset, and the fixation ring remained on the screen until 1.2 s after sentence offset. In Experiment 1, a 30-talker babble noise was presented throughout; that is, from three seconds prior to sentence onset until 1.2 s post-sentence offset (Figure 1A). In Experiment 2, no sound stimulation was administered during the three seconds prior to sentence onset and during the 1.2-s post-sentence offset period. A sentence started playing three seconds after the onset of the fixation ring (and 30-talker babble in Experiment 1). To ensure that participants tried to comprehend each sentence, and to assess comprehension, a semantic relatedness judgement was required after each sentence. The fixation ring on the screen was replaced by a probe word (e.g., “Book”) 1.2 s after sentence offset. Participants had to indicate with a keypress whether the probe word was semantically related or unrelated to the sentence they had heard. The word remained on screen for 3.5 seconds or until participants pressed the related (left index finger) or unrelated (right index finger) button on a keyboard, whichever came first. The screen was cleared between trials for 5-7 seconds in order to allow participants to rest and blink. Participants were instructed to maintain fixation and reduce blinks as long as the fixation ring was presented on the screen (including during presentation of sound materials).

Before both experiments, participants underwent a training block of 8 trials (using sentences not used in the experiment) in order to familiarize them with the experimental procedures (including eyetracker calibration). The experiment took approximately one hour to complete.

### Data analysis

Data analysis was carried out offline using custom MATLAB scripts (v2018b), and the analyses were identical for both experiments.

#### Behavior

The semantic relatedness responses were analyzed by calculating the proportion of correct responses, separately for each ambiguity and speech-clarity condition. A correct response entailed responding with “related” when a word was semantically related to the preceding sentence or by pressing “unrelated” when the word was not semantically related to the preceding sentence. Separately for each experiment, a 2 × 2 repeated-measures analysis of variance (rmANOVA) was calculated, with factors Clarity (Exp 1: +6 dB SNR, 0 dB SNR; SNR; Exp 2: clear, −2 dB SNR) and Ambiguity (LA, HA). Any significant interaction was resolved by subsequent t-tests.

#### Pupillometry

Preprocessing of pupil area involved removing eye-blink artifacts. For each eye blink indicated by the eye tracker, all data points between 50 ms before and 200 ms after a blink were removed. In addition, pupil area values that differed from the median pupil area by more than three times the median absolute deviation were classified as outliers and removed (Leys et al., 2013). Missing data resulting from artifact rejections and outlier removal were linearly interpolated. Data for an entire trial were excluded from analysis if missing data made up more than 40% of the trial. Data were low-pass filtered at 10 Hz (Kaiser window, length: 201 points). Single-trial time courses were baseline-corrected by subtracting the mean pupil size from the −0.5 s to 0 s time window from the pupil size value at each time point (Mathôt et al., 2018). Single-trial time courses were averaged separately for each condition, and displayed for the −0.5 s to 4 s epoch.

Three dependent measures were extracted: Mean pupil dilation, peak pupil dilation, and peak pupil latency (Winn et al., 2018). In order to account for the different sentence durations at the analysis stage, mean pupil dilation was calculated for each trial as the average pupil area within 0.5 s post sentence onset and 1 s post sentence offset, and subsequently averaged across trials, separately for each condition and participant. Peak dilation and peak latency were extracted for each trial within 0.5 s post sentence onset and 1 s post sentence offset, and subsequently averaged across trials, separately for each condition and participant.

Separately for each experiment and each dependent measure, a 2 × 2 rmANOVA was calculated, with factors Clarity (Exp 1: +6 dB SNR, 0 dB SNR; SNR; Exp 2: clear, −2 dB SNR) and Ambiguity (LA, HA). Any significant interaction was resolved by subsequent t-tests.

#### Microsaccades

Participants were instructed to maintain fixation and reduce blinks during a trial. Microsaccades commonly occur during prolonged fixation in auditory tasks (Widmann et al., 2014), as was used here, and microsaccades can influence pupil dilation (Knapen et al., 2016). We therefore tested the extent to which microsaccades show effects of speech clarity and semantic ambiguity. Microsaccades were identified using a method that computes thresholds based on velocity statistics from eyetracker data, and then identfies microsaccades as events passing that threshold (Engbert and Kliegl, 2003; Engbert, 2006). That is, the veritical and horizontal eye movement time series were transformed into velocities and microsaccades were classified as outliers if they exceeded a relative velocity threshold of 15 times the standard deviation of the eye-movement velocity and persisted for 6 ms or longer (Engbert and Kliegl, 2003; Engbert, 2006). A time course of microsaccade rate was calculated from the individual microsaccade times (Widmann et al., 2014) by convolving each microsaccade occurrence with a Gaussian window (standard deviation of 0.02 s; zero phase lag). Mean microsaccade rate was calculated across trials as the average rate in the time window ranging from 0.5 s post sentence onset to 1 s post sentences offset, and subsequently averaged across trials (similar to the analysis of mean pupil dilation). For display purposes, time courses of mean microsaccade rate were calculated for the −0.5 to 4 s time window relative to sentence onset.

Separately for each experiment, a 2 × 2 rmANOVA was calculated for the mean microsaccade rate, with factors Clarity (Exp 1: +6 dB SNR, 0 dB SNR; SNR; Exp 2: clear, −2 dB SNR) and Ambiguity (LA, HA). Any significant interaction was resolved by subsequent t-tests.

## Results

### Experiment 1

#### Semantic-relatedness task

Mean proportion correct in the semantic relatedness task exceeded 0.8, even in the most challenging condition (Figure 2). The rmANOVA on these data revealed that proportion correct was higher at +6 dB SNR than at 0 dB SNR (Clarity: F_1,37_ = 45.133, p = 6.74^e-8^, η^2^_*p*_ = 0.549). The main effect of Ambiguity was not significant (F_1,37_ = 3.148, p = 0.084, η^2^_*p*_ = 0.078), but the Clarity × Ambiguity interaction was significant (F_1,37_ = 8.118, p = 0.007, η^2^_*p*_ = 0.179), such that participants performed worse for high-ambiguity sentences compared to low-ambiguity sentences at 0 dB SNR (t_37_ = 3.03, p = 0.004), but not at +6 dB SNR (t_37_ = −0.544, p = 0.589).

**Figure 2.**
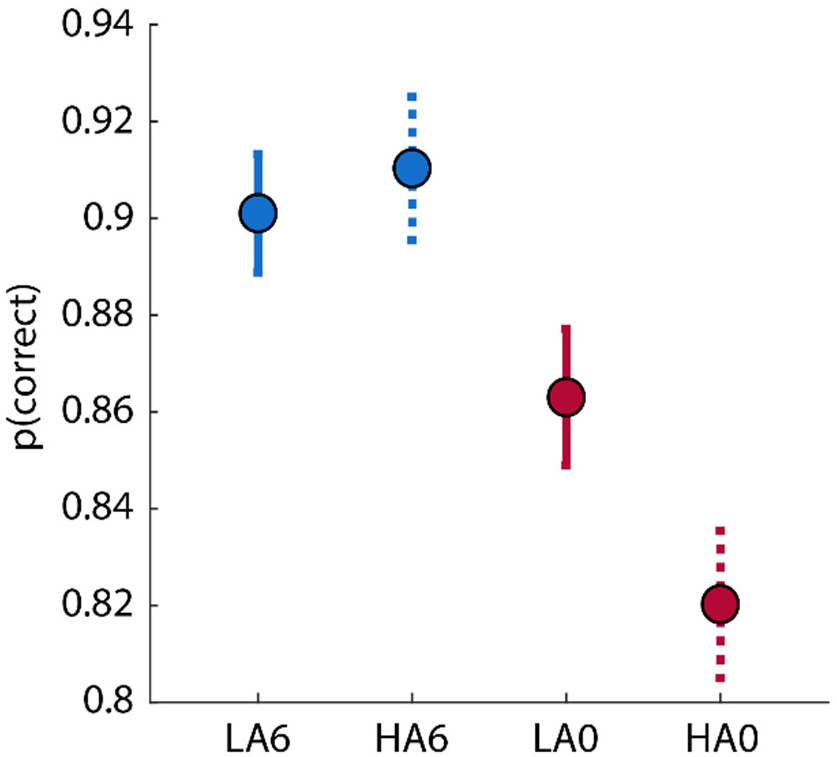
Experiment 1: Proportion correct in semantic-relatedness task. Mean proportion of correct responses for each condition. Error bars reflect the standard error of the mean. The Clarity × Ambiguity interaction was significant. LA6 – low ambiguity in +6 dB SNR babble, HA6 – high ambiguity in +6 dB SNR babble, LA0 – low ambiguity in 0 dB SNR babble, HA0 – high ambiguity in 0 dB SNR babble.

#### Pupillometry

Pupil area time courses are displayed in Figure 3A. The rmANOVA for the mean pupil area revealed that the pupil area was larger at 0 dB SNR than at +6 dB SNR (Clarity: F_1,37_ = 10.3, p = 0.002, η^2^_*p*_ = 0.218; Figure 3B,F). In addition, pupil area tended to be larger for HA sentences compared to LA sentences (marginally significant main effect of Ambiguity: F_1,37_ = 3.73, p = 0.061, η^2^_*p*_ = 0.091; Figure 3B,E). Individual data points are shown in Figures 3E and F. The Clarity × Ambiguity interaction also approached significance (F_1,37_ = 3.90, p = 0.055, η^2^_*p*_ = 0.095). Pupil area was larger in HA compared to LA sentences at +6 dB SNR (t_37_ = 2.953, p = 0.0054), but not at 0 dB SNR (t_37_ = 0.3552, p = 0.724).

**Figure 3.**
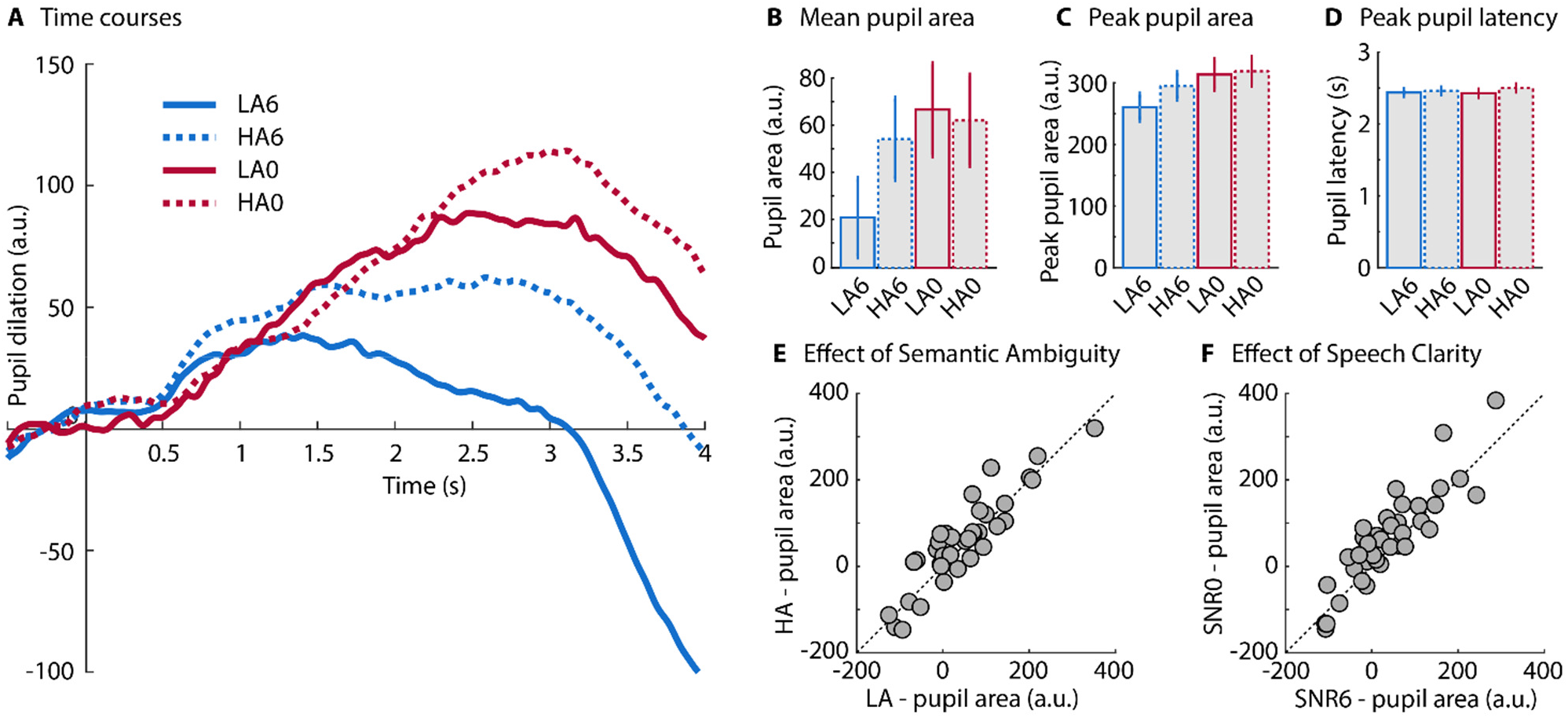
Experiment 1: Pupil dilation results. **A:** Time course of pupil area (averaged across participants; N=38) for sentences with durations between 2 and 3 s (74 out of 112 sentences), so as to minimize the impact of variance in sentence duration on the display of pupil time courses. **B:** Mean pupil area from 0.5 s after sentence onset to one second after sentence offset. **C:** Peak pupil dilation. **D:** Latency of peak pupil dilation. Error bars reflect the standard error of the mean. **E:** Individual data scatter plot for Ambiguity main effect (mean pupil area). **F:** Individual scatter plot for Clarity main effect (mean pupil area). LA6 – low ambiguity in +6 dB SNR babble, HA6 – high ambiguity in +6 dB SNR babble, LA0 – low ambiguity in 0 dB SNR babble, HA0 – high ambiguity in 0 dB SNR babble.

The rmANOVA for peak pupil area revealed that peak pupil dilation was larger at 0 dB SNR than at +6 dB SNR (Clarity: F_1,37_ = 18.1, p = 1.3^e-4^, η^2^_*p*_ = 0.328) and larger for HA compared to LA sentences (Ambiguity: F_1,37_ = 4.7212, p = 0.036, η^2^_*p*_ = 0.113; Figure 3C). The Clarity × Ambiguity interaction was not significant (F_1,37_ = 2.19, p = 0.1465).

The rmANOVA on peak latency revealed no significant main effects (Clarity: F_1,37_ = 0.26, p = 0.61; Ambiguity: F_1,37_ = 3.48, p = 0.069) and no interaction (F_1,37_ = 0.53, p = 0.468; Figure 3D).

In sum, Experiment 1 demonstrates that pupil area is sensitive to speech clarity and semantic ambiguity, indicating that both acoustic and linguistic factors affect pupil dilation. In both conditions, a babble noise was used as the masker, which may have introduced some informational masking, likely requiring cognitive control and distracter suppression (Johnsrude and Rodd, 2016), as well as energetic masking. In Experiment 2, we used a pink noise masker with an constant SNR of −2 dB relative to the spoken sentences: this masker was used to investigate whether pupil dilation is also sensitive to linguistic factors when energetic masking is more static, in which case some of the speech signal is obliterated and missing information must be inferred from the bits of speech that are perceived, likely requiring effective working memory and access to semantic knowledge (Johnsrude and Rodd, 2016).

### Experiment 2

#### Semantic-relatedness task

Proportion correct in the semantic relatedness task exceeded 0.85, even in the most challenging condition (Figure 4). The proportion of correct responses was lower for −2 dB SNR compared to clear sentences (Clarity: F_1,34_ = 22.69, p = 3.467^e-5^, η^2^_*p*_ = 0.4). The effect of Ambiguity was not significant (F_1,34_ = 0.879, p = 0.354, η^2^_*p*_ = 0.025), but a significant Clarity × Ambiguity interaction (F_1,34_ = 6.23, p = 0.017, η^2^_*p*_ = 0.155) was due to lower performance for HA compared to LA sentences at −2 dB SNR (t_34_ = −2.369, p = 0.023), but not for clear sentences (t_34_ = 1.74, p = 0.089).

**Figure 4.**
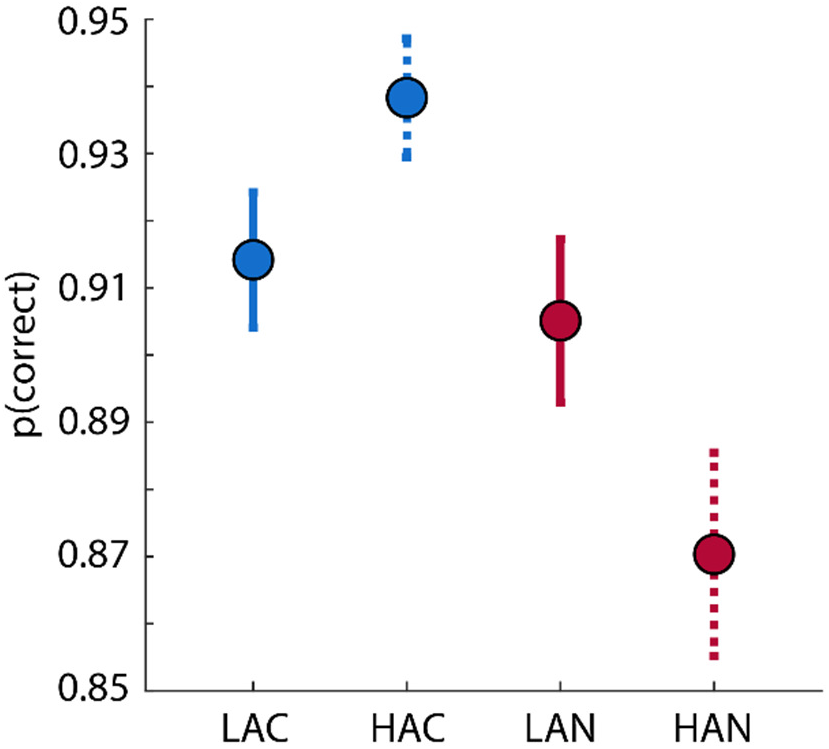
Experiment 2: Proportion correct in semantic-relatedness task. Mean proportion of correct responses for each condition. Error bars reflect the standard error of the mean. The Clarity × Ambiguity interaction was significant. LAC – low ambiguity in clear, HAC – high ambiguity in clear, LAN – Low ambiguity in −2 dB SNR pink noise, HAN – High ambiguity in −2 dB SNR pink noise.

#### Pupillometry

Pupil area time courses are displayed in Figure 5A. The rmANOVA for the mean pupil area revealed that mean pupil area was larger at −2 dB SNR compared to clear sentences (Clarity: F_1,34_ = 55.68, p = 1.169^e-8^, η^2^_*p*_ = 0.621; Figure 5B,F). Mean pupil area was also larger for HA than for LA sentences (Ambiguity: F_1,34_ = 5.53, p = 0.024, η^2^_*p*_ = 0.14; Figure 5B,E). Individual data points are shown in Figures 5E and F. The Clarity × Ambiguity interaction was not significant (F_1,34_ = 1.80, p = 0.188).

**Figure 5.**
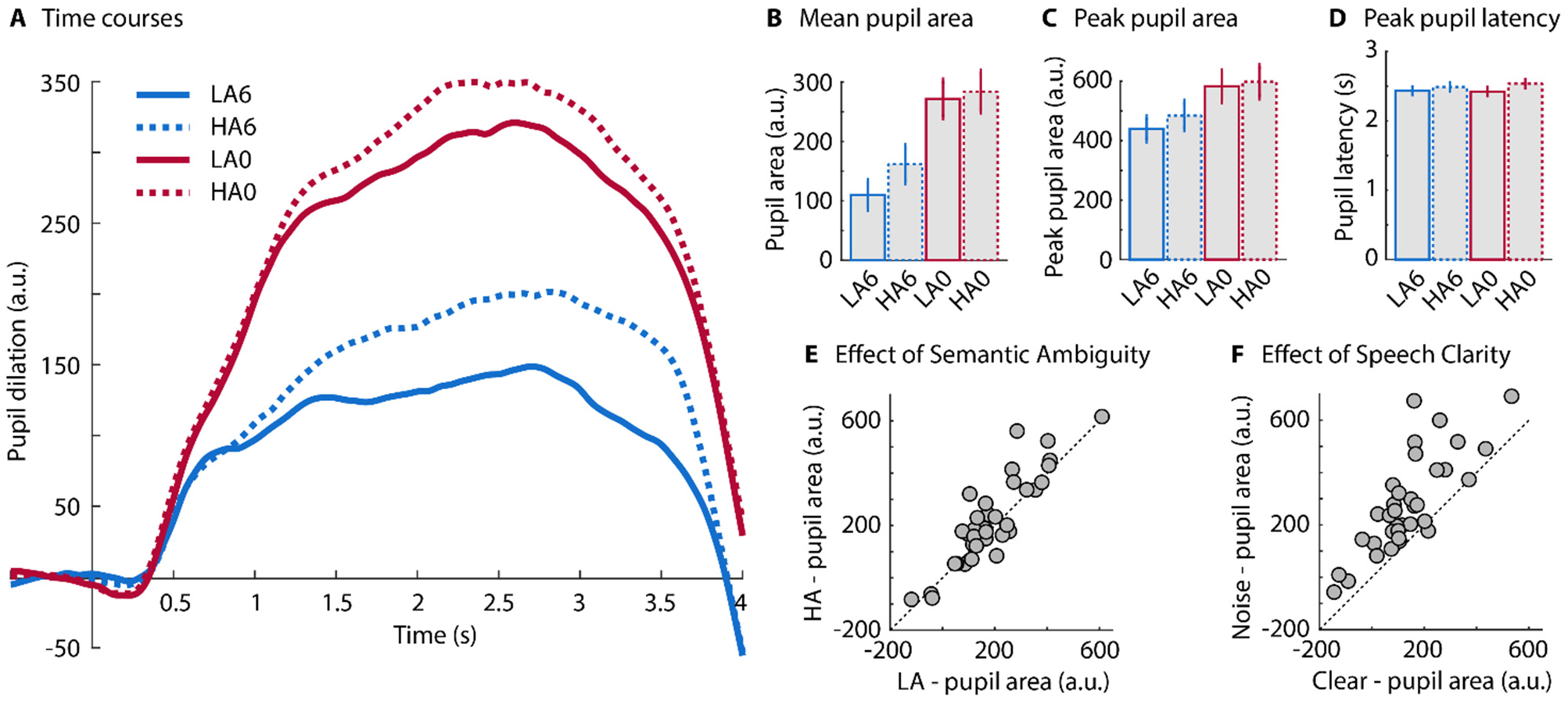
Experiment 2: Pupil dilation results. **A:** Time course of pupil area (averaged across participants; N=35) for sentences with durations between 2 and 3 s (74 out of 112 sentences), so as to minimize the impact of variance in sentence duration on the display of pupil time courses. **B:** Mean pupil area from 0.5 s after sentence onset to one second after sentence offset. **C:** Peak pupil dilation. **D:** Peak pupil latency. Error bars in B, C, and D reflect the standard error of the mean. **E:** Individual data scatter plot for the Ambiguity main effect (mean pupil area). **F:** Individual data scatter plot for Clarity main effect (mean pupil area). LAC – low ambiguity in clear, HAC – high ambiguity in clear, LAN – Low ambiguity in −2 dB SNR pink noise, HAN – high ambiguity in −2 dB SNR pink noise.

The results for peak pupil area mirrored those for mean pupil area (Figure 5C). Peak pupil area was larger in the noise compared to the clear conditions (Clarity: F_1,34_ = 53.57, p = 1.77^e-8^, η^2^_*p*_ = 0.611), and larger in the HA than in LA conditions (Ambiguity: F_1,34_ = 6.729, p = 0.0139, η^2^_*p*_ = 0.165). The Clarity × Ambiguity interaction was not significant (F_1,34_ = 0.2834, p = 0.283).

The rmANOVA for peak latency (Figure 5D) revealed that pupil size peaked later for HA than for LA sentences (Ambiguity: F_1,34_ = 11.48, p = 0.001). There was no effect for Clarity (F_1,34_ = 0.221, p = 0.640) and no Clarity × Ambiguity interaction (F_1,34_ = 1.8029, p = 0.188).

### Pooling data from Experiments 1 and 2

To compare behavioral performance in the semantic relatedness task across experiments, to gain more statistical power to observe any Clarity × Ambiguity interaction on pupil area, and to explore correlations between behavioral performance and pupil variables, we pooled the data from Experiments 1 and 2 (N=73). We performed rmANOVAs as before, with Experiment as a between-subjects factor.

#### Semantic relatedness task

Behavioral performance was higher in Experiment 2 compared to Experiment 1 (F_1,71_ = 5.095, p = 0.0271, η^2^_*p*_ = 0.067). Behavioral performance was higher for high compared to low SNR conditions (Clarity: F_1,71_ = 75.909, p < 0.00001, η^2^_*p*_ = 0.517). The main effect of Ambiguity was not significant (F_1,71_ = 2.972, p = 0.089), but there was a Clarity × Ambiguity interaction (F_1,71_ = 14.905, p = 0.000247, η^2^_*p*_ = 0.174) such that performance was lower for HA compared to LA sentences at low SNRs (t_72_ = 3.682, p = 0.0004) and trended towards higher performance on HA compared to LA sentences in high-SNR conditions (t_72_ = −1.941, p = 0.0562). The effect of Clarity was larger in Experiment 1 compared to Experiment 2 (Clarity × Experiment interaction: F_1,71_ = 4.701, p = 0.034, η^2^_*p*_ = 0.062), but not the effect of Ambiguity (Ambiguity × Experiment interaction: F_1,71_ = 0.798, p = 0.375).

#### Pupillometry

Mean pupil area was larger in Experiment 2 compared to Experiment 1 (F_1,71_ = 26.65, p = 2^e-6^, η^2^_*p*_ = 0.273). Mean pupil area was larger for low SNR compared to high SNR conditions (Clarity: F_1,71_ = 69.7, p = 3.7^e-12^, η^2^_*p*_ = 0.496) and larger for HA compared to LA sentences (Ambiguity: F_1,71_ = 9.32, p = 0.003, η^2^_*p*_ = 0.116). The Clarity × Ambiguity interaction (F_1,71_ = 4.97, p = 0.029, η^2^_*p*_ = 0.066) revealed that pupil area was larger for HA compared to LA sentences at high SNRs (t_72_ = 3.276, p = 0.002), but not at low SNRs (t_72_ = 0.351, p = 0.726). The effect of Clarity was larger in Experiment 2 compared to Experiment 1 (Clarity × Experiment interaction: F_1,71_ = 32.81, p < 0.00001, η^2^_*p*_ = 0.316), but not the effect of Ambiguity (Ambiguity × Experiment interaction: F_1,71_ = 1.395, p = 0.242).

The rmANOVA for peak pupil area mirrored the results for mean pupil area. Peak pupil area was larger in Experiment 2 compared to Experiment 1 (F_1,71_ = 22.349, p = 0.000011, η^2^_*p*_ = 0.239). Peak pupil area was larger for low SNR compared to high SNR conditions (Clarity: F_1,71_ = 74.859, p < 0.00001, η^2^_*p*_ = 0.513) and larger for HA compared to LA sentences (Ambiguity: F_1,17_ = 11.625, p = 0.0011, η^2^_*p*_ = 0.141). The Clarity × Ambiguity interaction approached significance (F_1,17_ = 3.161, p = 0.080, η^2^_*p*_ = 0.043), showing that pupil area was larger for HA compared to LA sentences at high SNRs (t_72_ = 3.608, p = 0.0006), but not for low SNRs (t_72_ = 0.934, p = 0.353). The effect of Clarity was larger in Experiment 2 compared to Experiment 1 (Clarity × Experiment interaction: F_1,71_ = 21.711, p = 0.000014, η^2^_*p*_ = 0.234), but not the effect of Ambiguity (Ambiguity × Experiment interaction: F_1,71_ = 0.479, p = 0.491).

#### Correlation between behavioral performance and pupil area

We examined whether comprehension (indexed by performance on the relatedness task) was related to pupil area by calculating correlations between behavioral performance and mean pupil area, partialing out Experiment so as to avoid biasing correlations by overall differences between experiments. No significant correlations were observed. The correlation between performance and pupil area, collapsed across clarity and ambiguity levels, was not significant (r = −0.218, p = 0.065, df = 70). The correlation between the HA vs. LA difference in behavioral performance and the HA vs LA difference in mean pupil area, collapsed across clarity levels, was also not significant (r = 0.197, p = 0.097, df = 70) and neither was the correlation between the low vs. high SNR difference in behavioral performance and low vs. high SNR difference in mean pupil areas, collapsed across ambiguity levels (r = 0.089, p = 0.455, df = 70). Finally, the correlation between the HA vs. LA difference in behavioral performance and the HA vs. LA difference in mean pupil area was not significant at high SNRs (r = 0.117, p = 0.330, df = 70) nor at low SNRs (r = −0.098, p = 0.414, df = 70). Thus, there appears to be no relation between mean pupil area and comprehension, at least as indexed by the semantic relatedness task used here.

### Microsaccade results

Microsaccades were analyzed in order to investigate whether saccadic eye movements during fixation are also sensitive to speech clarity and semantic ambiguity. Microsaccade time courses are depicted in Figure 6. The initial decrease in microsaccade rate after sentence onset is consistent with previous work showing a transient reduction in microsaccade rate for task-relevant auditory stimuli (Widmann et al., 2014).

**Figure 6.**
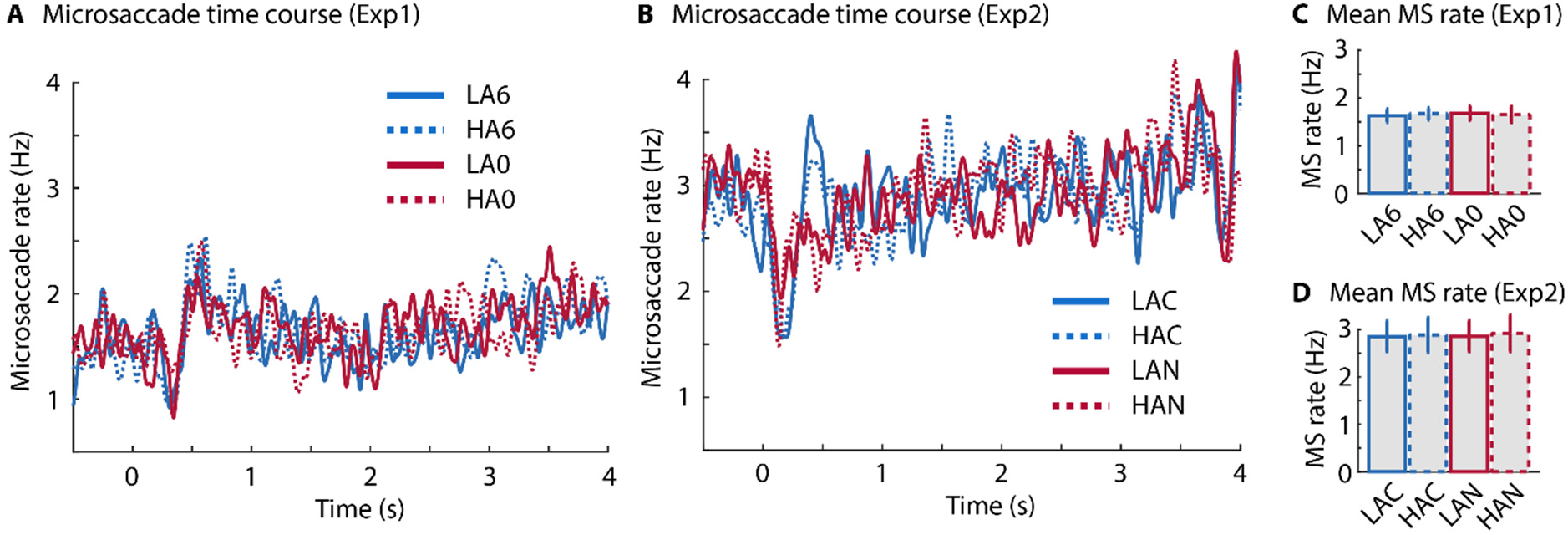
Results for microsaccade analysis. Time courses of microsaccade rate for Experiment 1 (A) and Experiment 2 (B). Bar graphs show the mean microsaccade rate for each condition for Experiment 1 (C) and Experiment 2 (D). Error bars reflect the standard error of the mean. MS – microsaccade. Exp 1 abbreviations: LA6 – low ambiguity in +6 dB SNR babble, HA6 – high ambiguity in +6 dB SNR babble, LA0 – low ambiguity in 0 dB SNR babble, HA0 – high ambiguity in 0 dB SNR babble. Exp 2 abbreviations: HAC – high ambiguity in clear, LAN – Low ambiguity in −2 dB SNR pink noise, HAN – High ambiguity in −2 dB SNR pink noise.

Microsaccade rate, averaged across the epoch spanning 0.5 s post-sentence onset to 1 s post-sentence offset, is shown in Figure 6C and D. No significant main effects or interactions were observed in the two experiments (Experiment 1: Clarity: F_1,37_ = 0.051, p = 0.821, η^2^_p_ = 0.001; Ambiguity: F_1,37_ = 0.003, p = 0.956, η^2^_p_ = 8^e-5^; Clarity × Ambiguity interaction: F_1,37_ = 0.316, p = 0.577, η^2^_p_ = 0.008; Experiment 2: Clarity: F_1,34_ = 0.039, p = 0.844, η^2^_p_ = 0.001; Ambiguity: F_1,34_ = 0.122, p = 0.729, η^2^_p_ = 0.003; Clarity × Ambiguity interaction: F_1,34_ = 0.017, p = 0.895, η^2^_p_ = 5^e-4^).

A rmANOVA conducted for microsaccade data pooled across experiments was conducted, with

Experiment as a between-subjects factor. The rmANOVA revealed a lower microsaccade rate in Experiment 1 compared to Experiment 2 (F_1,71_ = 12.08, p = 0.001, η^2^_*p*_ = 0.145).

## Discussion

### Speech comprehension

In the current study, we conducted two experiments to investigate the effects of speech clarity and semantic ambiguity on sentence comprehension and pupil dilation. Speech comprehension was good throughout as indexed by a semantic-relatedness task (all scores higher than 82% correct), but was reliably lower for acoustically degraded compared to less degraded sentences in both experiments, as expected (e.g., Miller, 1947; Cherry, 1953; Mattys et al., 2012; Johnsrude et al., 2013; Johnsrude and Rodd, 2016; Ohlenforst et al., 2017).

Comprehension was also lower for sentences containing homophones than for matched sentences without, but only at the lower SNRs (0 dB but not +6 dB in Experiment 1; and with noise but not clear in Experiment 2). This is interesting given that comprehension was still high, and that the low-ambiguity and high-ambiguity sentences are acoustically very similar. This effect may be due to the fact that contextual constraints are weaker in high-ambiguity compared to low-ambiguity sentences. Because we used meaningful sentences, their intelligibility (and thus performance on the comprehension task) is due to at least two factors. First, the acoustic quality of the signal determines intelligibility. Second, the sentence-level meaning (the context) imposes constraints that allows participants to ‘fill in’ the words they didn’t hear very well, using the words that they did. In low-ambiguity sentences, each of the content words has one meaning and these meanings can constrain interpretation. Listeners can use the words they perceive from acoustically degraded low-ambiguity sentences to generate a relatively small set of hypotheses regarding the identity of segments that they hear less well, and then ‘choose to hear’ words that fit with the overall meaning of the sentence. This process is less constrained for high-ambiguity sentences. Each homophone is semantically consistent with a wider set of hypotheses regarding the identity of less-well-heard sentence segments, and the overall meaning of the sentence is not as straightforward, since it depends on the constraints imposed mutually across all the homophones in the sentence (shell…fired…tank) and not on any one word perceived in isolation. Our observation of reduced comprehension by the presence of homophones is consistent with prior work indicating that homophones in naturalistic sentences introduce increased cognitive load (compared to matched sentences without homophones) as indexed by: 1) longer reaction times on a concurrent case-judgement task (Rodd et al., 2010a); 2) greater activity in functional MRI experiments (Rodd et al., 2005; Rodd et al., 2010b; Rodd et al., 2012; Rodd et al., 2015); and 3) poorer recognition memory (Koeritzer et al., 2018). Like Koeritzer et al., 2018 we also observe that the challenges introduced by homophony are particularly evident when the signal is of lower quality.

### Pupillometric measures

Pupil dilation, measured both as average area and peak area during sentence listening, was enhanced for acoustically degraded compared to less degraded sentences. This finding is in line with several previous observations demonstrating an enhanced pupil size when individuals listen under acoustic challenges (Zekveld et al., 2010; Koelewijn et al., 2014; Zekveld and Kramer, 2014; Winn et al., 2015; Wendt et al., 2016; Miles et al., 2017). Acoustic degradation due to auditory peripheral damage is associated with similar effects on pupil dilation during speech comprehension: It is larger for older compared to younger adults (Ayasse and Wingfield, 2018), for older adults with hearing loss compared to those without (Ayasse and Wingfield, 2018; but see Koelewijn et al., 2017; Wang et al., 2018), and for people with cochlear implants compared to people without (Winn, 2016).

Previous work and our findings suggest that different types of acoustic challenges all lead to enhanced pupil size. Degradation of the speech signal using noise vocoding (Winn, 2016), stationary noise (Zekveld et al., 2010; Ohlenforst et al., 2018), fluctuating noise (Koelewijn et al., 2014; Wendt et al., 2018), a single talker (Koelewijn et al., 2014; Wendt et al., 2018), multi-talker babble (Wendt et al., 2016; Ohlenforst et al., 2018; Wendt et al., 2018; current Figure 3), or noise correlated with a sentence’s amplitude envelope (current Figure 5), all increase pupil dilation relative to less-demanding control stimuli. However, just because the pupillary manifestation is similar across challenges does not mean that the cognitive resources being recruited are the same. As reviewed in the introduction, different demands probably recruit different processes (Johnsrude and Rodd, 2016). The pupil was larger when participants listened to everyday, naturalistic, sentences containing homophones compared to matched sentences without homophones. This is in line with the observation that pupil dilation increases for isolated words that are presented in the context of lexical competitors (Kuchinsky et al., 2013) or are otherwise semantically difficult to process (based on word frequency, familiarity, naming latency, and age of acquisition) (Kuchinke et al., 2007; Chapman and Hallowell, 2015) compared to control words. Moreover, sentences in which semantic context does not predict the sentence’s final word lead to larger pupil dilation compared to sentences with a final word more predicable from context (Winn, 2016). Other work has demonstrated that pupil dilation increases when individuals listen to syntactically complex sentences compared to less complex ones (Wendt et al., 2016; Ayasse and Wingfield, 2018; but see Müller et al., 2019). Consistent with Kahneman’s early assertion (Kahneman and Beatty, 1966; Kahneman, 1973) that anything involving mental effort increases pupil dilation, these previous observations and our data show that not just the quality of the speech signal, but the cognitive/linguistic demands of the speech signal increase pupil dilation. This is the case even when behavioral performance is unaffected (recall that comprehension performance did not differ between high-ambiguity and low-ambiguity sentences when these were presented clearly [Experiment 2] or at a higher SNR [Experiment 1]).

In addition to consistent main effects of clarity and ambiguity on pupil dilation, we further observed a consistent trend towards an interaction, with the difference in pupil response for high-ambiguity compared to low-ambiguity sentences being reliably larger when signal quality was better, compared to when it was poorer (Figures 3 and 5). The sub-additive effects of acoustic and linguistic challenges on pupil dilation is consistent with the suggestion that the pupil dilation saturates for highly degraded, but still-intelligible speech (Zekveld et al., 2014; Ohlenforst et al., 2017; Ohlenforst et al., 2018). Yet, a different interaction pattern has been observation in other work: Pupil dilation was increased when acoustic and linguistic challenges were present concurrently (compared to auditory stimuli without these challenges), but not for acoustic or linguistic challenges alone (Kuchinsky et al., 2013; late time window in Wendt et al., 2016). These data rather suggest super-additivity. Critically, the pupil area in the current study may have approached a physiological saturation, such that, in fact, the different cognitive processes recruited to compensate for degraded speech, and to cope with the presence of homophones, have an additive effect on the pupil.

### Relation between behavioral performance and pupil dilation

Comprehension behavior and pupil dilation appear to provide different windows on speech processing. At higher levels of clarity (+6 dB SNR in Experiment 1; clear presentation in Experiment 2) behavioral performance did not differ between high-ambiguity and low-ambiguity sentences, whereas pupil area was larger for high-ambiguity compared to low-ambiguity sentences. In contrast, at lower levels of clarity (0 dB SNR babble in Experiment 1, −2 dB SNR pink noise in Experiment 2) comprehension was reduced for high-ambiguity compared to low-ambiguity sentences, but the effect on pupil was not significant. These two apparently contradictory patterns may, to some extent, reflect saturation of the pupil, as discussed above.

In general, however, comprehension did not appear to relate to pupil area, even when saturation was not a factor. For example, comprehension was generally lower in Experiment 1 compared to Experiment 2, but the absolute magnitude of the pupil area (relative to pre-sentence baseline), indexing challenges/effort, was also smaller in Experiment 1 than in Experiment 2. Furthermore, the effect of clarity level on comprehension was larger in Experiment 1 (+6 dB vs 0 dB SNR in babble) than in Experiment 2 (clear vs −2 dB SNR pink noise), but the effect of clarity level on pupil dilation was smaller in Experiment 1 than in Experiment 2.

Although pupillometry recordings are increasingly used as a measure of listening effort (Winn et al., 2018; Zekveld et al., 2018), our data complement other data indicating that pupillometric measures do not always correlate with task performance measures or other measures of listening effort, such as subjective ratings or oscillatory neural activity (Hicks and Tharpe, 2002; Zekveld et al., 2010; Mackersie and Cones, 2011; Winn et al., 2015; Miles et al., 2017; Strand et al., 2018; Alhanbali et al., 2019). Part of the inconsistency may be due to the fact that the term ‘listening effort’ is ambiguous (Herrmann and Johnsrude, 2019) because it may refer to a mental act – associated with the recruitment of resources (Pichora-Fuller et al., 2016; Peelle, 2018) – or to a subjective experience (Johnsrude and Rodd, 2016; Lemke and Besser, 2016; Herrmann and Johnsrude, 2019). Motivation is another crucial factor, in addition to resource recruitment and listening experience, that determines how individuals engage in listening (Pichora-Fuller et al., 2016; Richter, 2016; Herrmann and Johnsrude, 2019). Different measures most certainly differ in the extent to which they tap into resource recruitment, motivation, and/or experience, making the absence of correlations between behavioral performance measures and physiological measures, as well as the absence of correlations among physiological measures, less surprising.

### Microsaccades are not influenced by semantic ambiguity and speech clarity

In the current experiments, participants were instructed to maintain fixation and reduce blinks during a trial. Microsaccades commonly occur during fixation (Engbert, 2006; Martinez-Conde et al., 2013; Widmann et al., 2014) and can influence pupil dilation (Knapen et al., 2016). Hence, microsaccades could in principle be entangled with changes in pupil size.

Here, we observed a transient inhibition in microsaccade rate following sentence onset (Figure 6). This is in line with previous observations that the probability of microsaccades is reduced following the onset of task-relevant auditory and visual stimuli (Rolfs et al., 2005; Rolfs et al., 2008; Widmann et al., 2014). Microsaccade inhibition is typically followed by an overshoot and a return to baseline (Rolfs et al., 2008; see also Figure 6). Critically, neither signal quality (clarity factor) nor the presence of homophones (ambiguity factor) affected microsaccade rate. The changes in pupil dilation induced by speech clarity and semantic ambiguity are therefore probably not related to microsaccades.

Analysis of microsaccade differences between experiments shows that the microsaccade rate was overall lower in Experiment 1 compared to Experiment 2 (Figure 6). Microsaccade rate has been shown to decrease with high cognitive load (Dalmaso et al., 2017; Xue et al., 2017) and task difficulty (Siegenthaler et al., 2014). This is in line with the overall lower performance in Experiment 1 compared to Experiment 2 but is in contrast to the overall larger pupil size (relative to baseline) and larger effect of speech clarity in Experiment 2 compared to Experiment 1. The contrasting effects observed in the current study are consistent with the observation that different measures of listening effort and cognitive load are not or only minimally correlated (Miles et al., 2017; Alhanbali et al., 2019).

## Conclusions

The current study investigated the effects of acoustic degradation and semantic ambiguity on sentence comprehension and pupil dilation. Sentences comprehension, as indexed by performance on a semantic-relatedness task, was generally high but was reduced by acoustic degradation and by semantic ambiguity. Pupil dilation increased when signal quality was poor, and when homophones were present in everyday, naturalistic sentences. These two effects appeared additive at least within the physiological limits of the pupil response, both when babble was used as a masker, and when pink noise was used as a masker. The current results reinforce the idea that many different challenges to speech comprehension, that afford different cognitive processes and are met by the brain in different ways, manifest as an increase in pupil dilation. When using pupillometry to measure listening effort specifically, other forms of “mental effort”, such as linguistic and domain-general abilities required to comprehend speech, and recruited only insofar as the speech signal requires them, must be controlled.

## Author contributions

MK helped design the study, collected data, analyzed data, and wrote the manuscript. BH designed and programmed the study, assisted with data analysis, and wrote the manuscript. JR provided the stimuli and edited the manuscript. ISJ conceived and designed the study and wrote the manuscript.

## Acknowledgements

This work was supported by NSERC and CIHR operating grants awarded to ISJ. BH was supported by a BrainsCAN Tier I postdoctoral fellowship (Canada First Research Excellence Fund; CFREF).

